# Early root phenotyping in sweetpotato (*Ipomoea batatas* L.) uncovers insights into root system architecture variability

**DOI:** 10.1101/2023.01.18.524552

**Authors:** Luis O. Duque

## Abstract

**Background:** We developed a novel, non-destructive, expandable, ebb and flow soilless phenotyping system to deliver a capable way to study early root system architectural traits in stem derived adventitious roots of sweetpotato (*Ipomoea batatas* L.). The platform was designed to accommodate up to 12 stems in a relatively small area for root screening. This platform was designed with inexpensive materials and equipped with an automatic watering system.

**Methods:** To test this platform, we designed a screening experiment for root traits using two contrasting sweetpotato genotypes, ‘Covington’ and ‘NC10-275’. We monitored and imaged root growth, architecture, and branching patterns every five days up to 20 days.

**Results:** We observed significant differences in both architectural and morphological root traits for both genotypes tested. After 10 days, root length, surface root area, and root volume were higher in ‘NC10-275’ compared to ‘Covington’. However, average root diameter and root branching density were higher in ‘Covington’.

**Conclusion:** These results validated the effective and efficient use of this novel root phenotyping platforming for screening root traits in early stem-derived adventitious roots. This platform allowed for monitoring and 2D imaging root growth over time with minimal disturbance and no destructive root sampling. This platform can be easily tailored for abiotic stress experiments, permit root growth mapping and temporal and dynamic root measurements of primary and secondary adventitious roots. This phenotyping platform can be a suitable tool for examining root system architecture and traits of clonally propagated material for a large set of replicates in a relatively small space.

**Subjects:** Plant Science, Agricultural Science

## Introduction

Crop productivity is mainly influenced by environmental factors which include, but are not limited to fluctuating temperatures, seasonal radiant energy, available and accessible soil moisture and mineral nutrients that are distributed in soil. Of the abovementioned factors, soil moisture and mineral nutrients directly affect growth and distribution of root systems (Purushothaman *et al*., 2017b; Purushothaman *et al*., 2017a; Siddique *et al*., 2015; Gao and Lynch, 2016; Burridge *et al*., 2016; Zhan and Lynch, 2015). Examining *in situ* or *ex situ* root systems are key to understanding crop productivity, as soil resources are heterogeneously dispersed in soil profiles or are prone to localized depletion, making root spatial growth and distribution shape the capacity of a plant to capitalize on available resources (Lynch, 1995). Research on improving root system architecture (RSA) under low nutrient, low input agriculture and water stress could improve overall crop yield (Wasson *et al*., 2012; Kuijken *et al*., 2015) and favorable changes in root architecture for nutrient capture and utilization of soil moisture could influence overall biomass accumulation, hence, yield (Hammer *et al*., 2009; Anami *et al*., 2015; Xie *et al*., 2017). However, the exploration of RSA traits is laborious due to the impediment of accessing the soil matrix.

To circumvent this constraint, numerous real-time growth monitoring systems have been developed for root visualization and quantification with support of innovative optical recording techniques used in greenhouse settings. Furthermore, newer methodologies and improvements on existing platforms for phenotyping large number of genotypes, replicates and treatments are being developed with more reliable results (Kuijken *et al*., 2015; Chen *et al*., 2011). These *exsitu* root phenotyping platforms can be categorized into two broad groups: 1) soil/substrate systems and 2) non-soil systems. The soil/substrate system consists of a rhizotron/-box/-mesocosm containing sand, natural soil or artificial soil mix were root growth is either monitored non-invasively with the use of X-ray micro-tomography, magnetic resonance imaging or CT scanning or destructively by digging, removing and cleaning whole root system and afterward, scanning and/or taking a picture for further analysis (Nagel *et al*., 2012; Blossfeld *et al*., 2011; Rascher *et al*., 2011; Metzner *et al*., 2015; Pflugfelder *et al*., 2017; Saengwilai *et al*., 2014; Zhan *et al*., 2015). The soilless systems include: hydroponics (Clark *et al*., 2013; Pace *et al*., 2014), agar or gellan gum (Fang *et al*., 2009; Iyer-Pascuzzi *et al*., 2010; Clark *et al*., 2011; Topp *et al*., 2013; Ribeiro *et al*., 2014), aeroponics (de Dorlodot *et al*., 2007; Gaudin *et al*., 2011), grow pouches (Hund *et al*., 2009; Adu *et al*., 2014), transparent soil (Downie *et al*., 2012) and rhizoslides (Le Marie *et al*., 2016). Both *ex situ* systems present advantages as well as disadvantages dependent on end results. For example, environmental unpredictability can be reduced using standardized artificial media, nutrient composition and/or application and micro-environment control of both systems. In addition, they have the capability of real-time direct root growth observations avoiding destructive harvest and can be very high throughput. On the other hand, the 2D or 3D nature of both systems force root growth and development in an unnatural physical realm as well as in a chemically artificial media. Lastly, both systems have the limitation of using seed and seedling growth as proxies for mature plants. Thus, the optimal phenotyping platform should accommodate a range of desirable properties, such as, low operating and developmental costs and the possibility of measuring large number of plants, replicates and/or treatments (Kuijken *et al*., 2015). Up to now, the majority of techniques developed for RSA phenotyping rely on the use of seedlings and early stage root phenotypes which have shown some predictive value for later developmental stages (Tuberosa *et al*., 2002), however other studies have shown that this is not the case when seedling root phenotypes are compared to a mature plant (Watt *et al*., 2013), hence, a flexible phenotyping system that would allow a time-series capture of several developmental stages would be of paramount importance and have increased agronomic relevance. Now, there is a lack of a suitable root phenotyping method enabling the study of time-series stem derived storage root systems for that is inexpensive, scalable, and adoptable by low resource or fund-limited laboratories worldwide. Here, we focused on the root system of the storage root crop, sweetpotato (*Ipomoea batatas* L.), an important and emerging crop for both developing as well as for developed nations worldwide.

Sweetpotato, is a vegetatively propagated true root crop that provides food security for resource-poor small holder farmers in Sub-Saharan Africa as well as in other tropical and sub-tropical countries worldwide (Khan *et al*., 2016). Limited literature is available on sweetpotato root growth and development when compared to cereals, and what is available, focuses on storage root growth, bulking and yield leaving out RSA entirely. Sweetpotato roots are adventitious roots (AR) originating from the shoot or underground stem (Khan *et al*., 2016), contrary to the root systems of seed propagated crops which consist of embryonic primary roots, seminal roots and stem borne crown roots (Hochholdinger *et al*., 2004; Hund *et al*., 2011). Sweetpotato RSA is composed of AR, lateral roots (LR) and storage roots (SR). The simple identification of a main AR axis and emerging LR through spatial and temporal events would enable novel research to further recognize mechanisms involved in LR emergence and function(Khan *et al*., 2016). To exploit early sweetpotato root traits as potential selection criteria for breeding programs that target different environmental scenarios, attempts have to be guided towards the development of 1) a robust and reproducible root phenotyping platform, 2) sustain stem and root growth until storage initiation, 3) express high heritability and/or repeatability for a given trait, 3) minimize genotype x environment interaction, 4) be able to be used all year around, and 5) not be labor intensive.

In our study, first we describe a novel, non-destructive, expandable, ebb and flow soilless phenotyping platform that is equipped with a customized imaging setup for stem derived (i.e., ‘slips’) storage root systems using germination paper that is preferred for large scale root phenotypic screens. And second, we examine the inherent genetic variations in root traits among a commercially available sweetpotato clone and an unreleased breeder line. This system enables the analysis of large number of replicates with relatively low-cost materials, non-destructive realtime direct root growth observations and imaging based on RGB photography and WinRhizo root image analysis.

## Materials & Methods

### Root phenotyping platform

Each individual phenotyping system consisted of a 17-gallon (64.3 liter) heavy duty polypropylene tough tote (26.88 in. L x 18 in. W x 12.5 in. H, HDX Model# SH17GTOUGHTLDBY, Home Depot, Atlanta, GA, U.S.A.) protected with a radiant barrier with a reflectance (IR) estimated at 94%+ (Reflectix Insulation, Markleville, IN, U.S.A.) (Figure 1A). The radiant barrier was used to prevent the phenotyping system from overheating caused by direct natural and artificial light. The original plastic top lid was removed and a retrofitted polyisocyanurate rigid foam insulation (Rmax Thermasheath-3, Dallas, TX, U.S.A.) was used in its place. The retrofitted foam insulation lid was cut with 12 rectangular openings (13 in. L x 1 in. W spaced 1 in. apart from opening to opening) to accommodate each individual growth unit (Figure 1B). Figure 2 shows a schematic representation of each phenotyping individual system.

**Figure 1.**
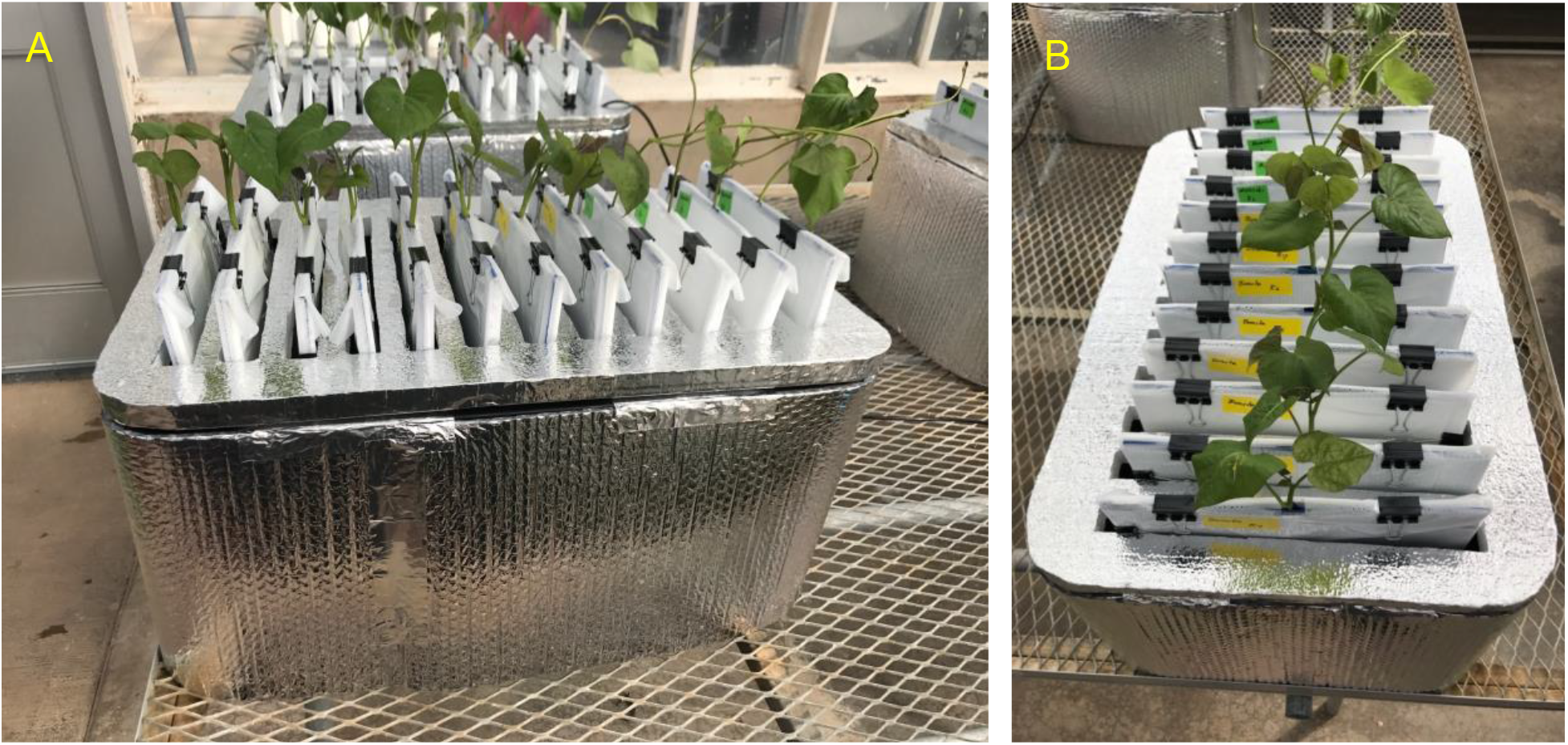
Sweetpotato ebb and flow soilless phenotyping platform constructed and tested for stem-derived adventitious roots: (A) side and (B) aerial view showing the 12 individual growth units fitted with sweetpotato ‘slips’.

**Figure 2.**
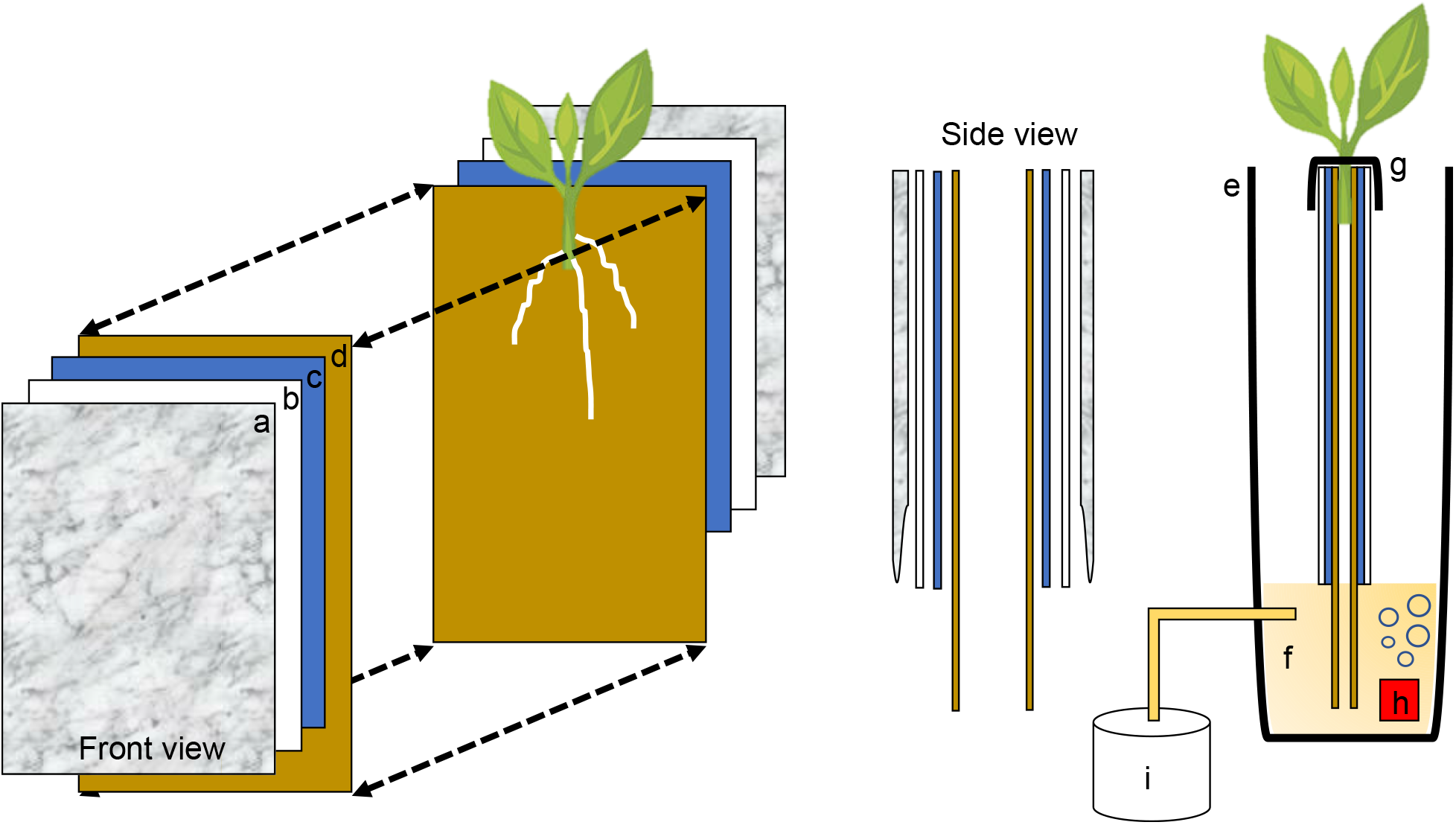
Schematic representation of the semi-hydroponic ebb and flow phenotyping system: (a) transparent plastic sheeting, (b) corrugated white plastic, (c) anchor steel blue seed germination paper, (d), brown heavy weight germination paper, (e) tank or reservoir (not drawn to scale), (f), nutrient solution, (g) securing clip, (h) bubbler, (i) nutrient tank retrofitted with an automatic submersible pump through a time controller.

### Individual growth unit

Each plant growth unit consisted of two sheets of heavy weight germination paper (18 in. H x 12 in. L, 76 lb., Anchor Paper Company, Saint Paul, MN, U.S.A.) followed by two sheets of Steel Blue Seed germination paper (18 in. H x 12 in. L; 120 lb.; Anchor Paper Company, Saint Paul, MN, U.S.A.) (Figure 3A). The germination paper was then sandwiched between two 0.157 in. thick white corrugated plastic sheets (15 in. H x 12 in. L, Coroplast Inc. Chicago, IL, U.S.A.) (Figure 3B) and then covered with 6 mm clear recycled polyethethylene sheeting (15 in. H x 12 in. L; HDX, Home Depot, Atlanta, GA). Four 1 ¼ in. metal binder clips (Staples, Framingham, MA, U.S.A.) were used to attached and hold together the clear plastic sheeting around the white corrugated plastic sheets. All germination paper was autoclaved (120 ^°^C for 20 min) and plastic sheeting was surface sterilized with 70% sodium hypochlorite and rinsed in deionized water.

**Figure 3.**
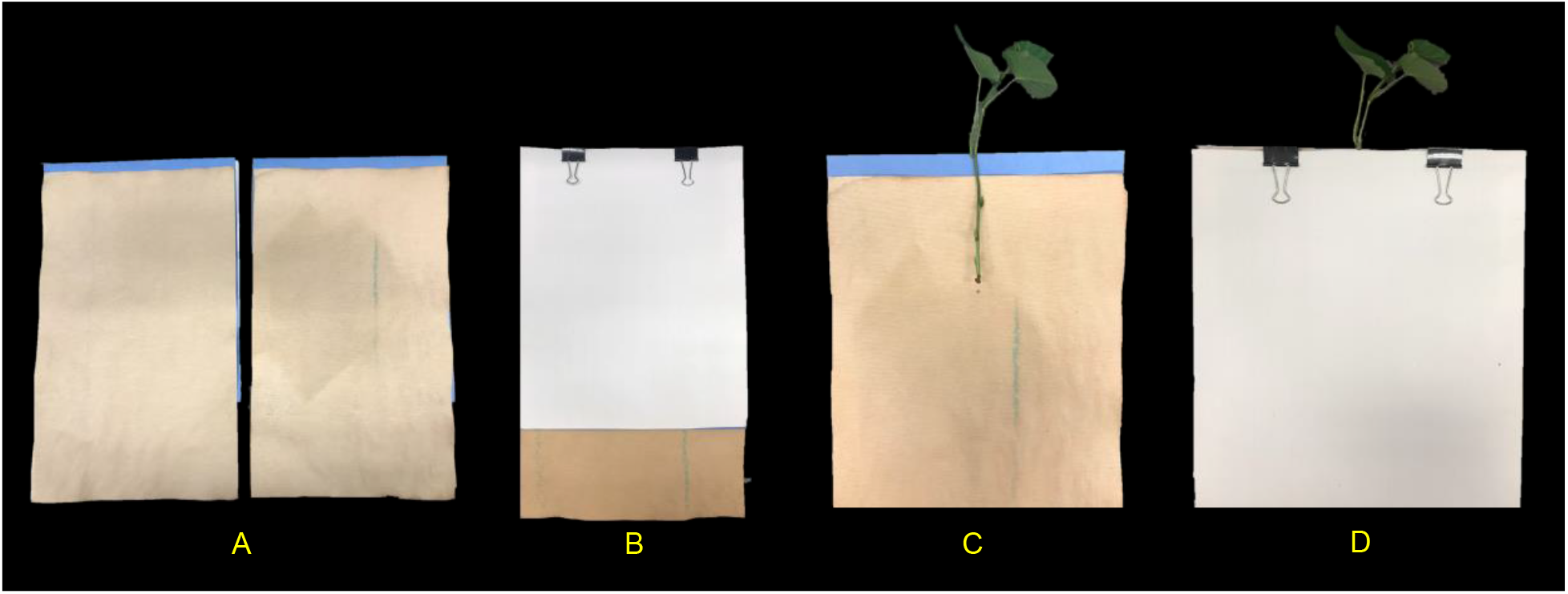
Individual growth units and ‘slip’ placement inside the growth unit: (A) left and right side of the growth unit showing germination paper placement before ‘slip’ positioning, (B) closed and “sandwiched” growth unit and secured with two metal binder clips (without ‘slip’), (C) ‘slip’ placement in the middle of one side of the growth unit and, (D) completed growth unit showing ‘slip’ protruding outward.

### Growth unit assembly and stem (‘slip’) placement in growth unit

With the rigid foam insulation lid covering the phenotyping system, each individual growth unit with one protruding stem was positioned into one rectangular opening and secured from the top with two 1 ¼ in. metal binder clips on each side (12 growth units per phenotyping system box). One excised sweetpotato stem was used per individual growth unit (Figure 3C). In short, stems from each genotype tested were randomly chosen and all mature leaves and petioles were excised leaving only two to three small immature leaves at the top. Each stem was them cut to a uniform length and care was taken so that two to three nodes were exposed and placed centered into the growth unit (pre-moisten with nutrient solution) with the rest of the stem with leaves protruding out (Figure 3D). Each growth unit was then closed and fastened with the metal binder clips and hung through one rectangular slot of the phenotyping platform.

### Irrigation System (Ebb-and-Flow System)

One 55-gallon heavy duty polypropylene tough tote with lid (45.43 in. L x 21.13 in. W x 19.52 in. H. HDX Model # HDX55GONLINE(4), Home Depot, Atlanta, GA, U.S.A.) was used as a nutrient storage tank. The tank was equipped with one 1/12 HP submersible pump (Model: Little Giant 4E-34NR Series; Franklin Electric Co., Inc., Fort Wayne, IN, U.S.A.). The pump was connected to a flexible ½ in. male national pipe thread hose (MNPT hose). The hose was then connected via an Ebb-and-Flow fitting kit (HydroFlow Products, Hawthorne Gardening Co., Vancouver, WA, U.S.A.) to one root phenotyping system. A digital timer was connected to the pump system for periodic water supply. The nutrient solution consisted of: 7 mM N, 0.5 mM P_2_O_5_; 7.5 mM K_2_O; 2 mM Mg; 2 mM S; 50 µM B; 10 µM Mn; 5 µM Zn; 2 µM Cu; 1 µM Mo. The nutrient solution stored in the nutrient supply tank was delivered to the phenotyping platform via an automatic submersible pump through a time controller. The periodic pumping was set as 10 min on and 240 min off during a 24-hour period. The nutrient solution was refreshed weekly.

### Location, genotypes, and growth condition

This study was conducted twice from September to October 2017 and from February to March 2018 in a temperature-controlled greenhouse at Penn State University located in University Park, PA, USA (40^°^48′N, 77^°^51′W). Greenhouse environmental growth conditions exhibited a photoperiod of 14/10 h at 32/28 °C (light/darkness) with a maximum midday photosynthetic flux density of 1200 *µ*mol photons m^−2^ supplemented with LED lights. The ambient humidity was 40%. One commercial and commonly available sweetpotato clone, ‘Covington’, and one unreleased breeder line, ‘NC10-275’, were tested throughout the system establishment of this root phenotyping platform.

### Data collection

Root growth was monitored, photographed, and measured every five days for a total of 20 days. With care, each growth unit was removed from the phenotyping platform and opened by removing the polyethethylene sheeting and white corrugated plastic sheets. Each growth unit was placed centered inside a light tent with built-in LED lights (Angler) and photographed with a standard DLSR camera (Canon; image size: 4000 × 6000 pixels; image DPI: 70 pixels/inch; color model: RGB; file type: JPEG) positioned on an adjustable overhead camera platform (Glide Gear) (Figure 4A and 4B). To account for root image scaling, a ruler was placed alongside each root.

**Figure 4.**
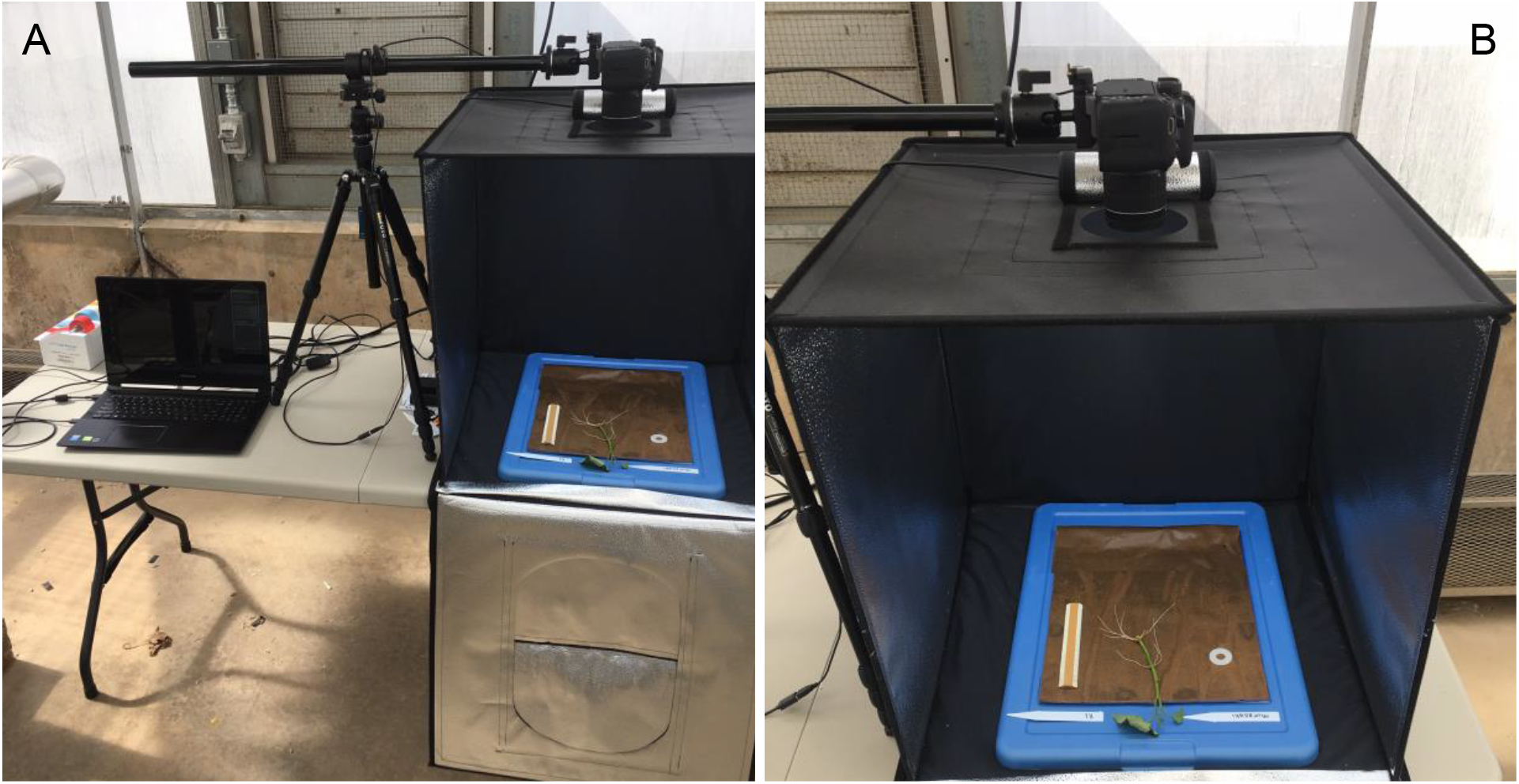
Imaging and data acquisition platform: (A) PC laptop connected to a standard DLSR digital camera positioned on an adjustable platform, and (B) light tent with built in LED lights showing the detail of the opened growth unit and exposing the ‘slip’ and root growth after 5 days.

### Image Analysis

All root images were pre-processed using Preview (Version 10.0; Apple Inc. Cupertino, CA, U.S.A.) and FIJI (Version 2.0.0-rc-69/1.52i; LOCI; University of Wisconsin-Madison, WI, U.S.A.). Preview was used to crop and remove the stem (i.e., stem/slip) segment from the rest of the root system using the *instant alpha* tool (Figure 5). This process was done manually to all root images. Cropped images were loaded to FIJI and 32-bit RGB (red, green, blue) images were converted to 8-bit grayscale LUT (look-up-table) for further processing. FIJI’s *subtract background* and *threshold* commands were then applied to separate roots from background (Figure 5).

**Figure 5.**
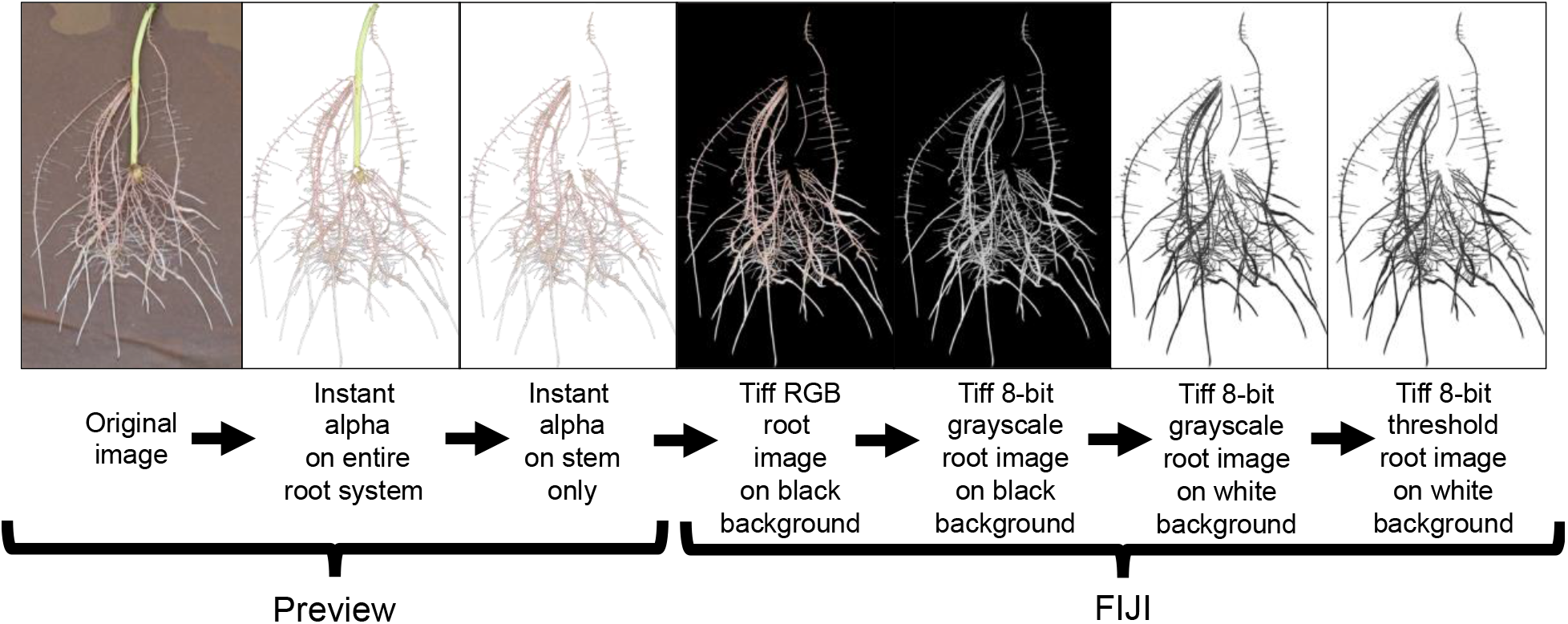
Root image processing sequence for subsequent root analysis in WinRhizo. This image preparation method includes a cropping of the background germination paper and stem using the Instant Alpha tool in Preview and then transferring the processed image to FIJI for image conversion and thresholding.

### Root Image Descriptors

For root morphological descriptors, WinRHIZO (V.2009 Pro, Regent Instruments, Montreal, QC, Canada) was used to detect root structures from each image. The diameter classes were set at 200 µm, the equivalent of two pixel with 10 equal intervals. The debris removal filter of WinRHIZO was set to remove objects with an area smaller than 0.02 cm^2^ and a length:width ratio lower than 10. WinRHIZO was able to distinguish adventitious/nodal and lateral roots in all images analyzed. All root morphological traits measured are listed in Table 1.

**Table 1.** Description of 10 measured traits for both sweetpotato genotypes grown in the root phenotyping platform evaluated at 5, 10, 15 and 20 days after placement in the growth unit.

| Trait Name | Unit | Trait Description |
| --- | --- | --- |
| Total Root Length | cm | Cumulative length of all roots in entire root system |
| Total Root Surface Area | $\text{cm}^2$ | Cumulative surface area of all roots in entire root system |
| Average Root Diameter | mm | Average diameter of all roots in entire root system |
| Total Root Volume | $\text{cm}^3$ | Cumulative volume of all roots in entire root system |
| Number of Root Tips | count | Cumulative tips of all roots in entire root system |
| Number of Root Forks | count | Cumulative forks of all roots in entire root system |
| Root Length Pattern | cm | Root length by depth at each 10 cm depth |
| Root Depth Index | % | Percent vertical centroid of root distribution in the soil |
| Root Branching Density | branch $\text{cm}^{-1}$ | Number of lateral branches per length unit along a root |
| Root Length Distribution by Diameter Class | cm | Cumulative root length by root diameter [very fine (0-0.6 mm), fine (>0.6 to $\leq 1.2$ mm), large (>1.2 to $\leq 1.8$ mm), and very large (>1.8 mm)] |

### Statistical Analysis

Analysis of variance was performed using JMP Pro 16 (SAS Institute, Cary, NC) on all measured root traits. Data were transformed, when necessary, before ANOVA to achieve normality. Tukey’s Honest Significant Difference test was used to identify significant differences among means. Statistical significance was based on a *p* value of < 0.05.

## Results

### Root development in the system

The root development for both genotypes tested were vigorous and presented root phenotype variation within the phenotyping system. The root systems of both ‘Covington’ and ‘NC10-275’ consisted of several first-order adventitious roots (i.e., lateral roots) originating from nodes placed with the phenotyping system and by the end of the experiment, second-order root branching was also observed (Figure 6).

**Figure 6.**
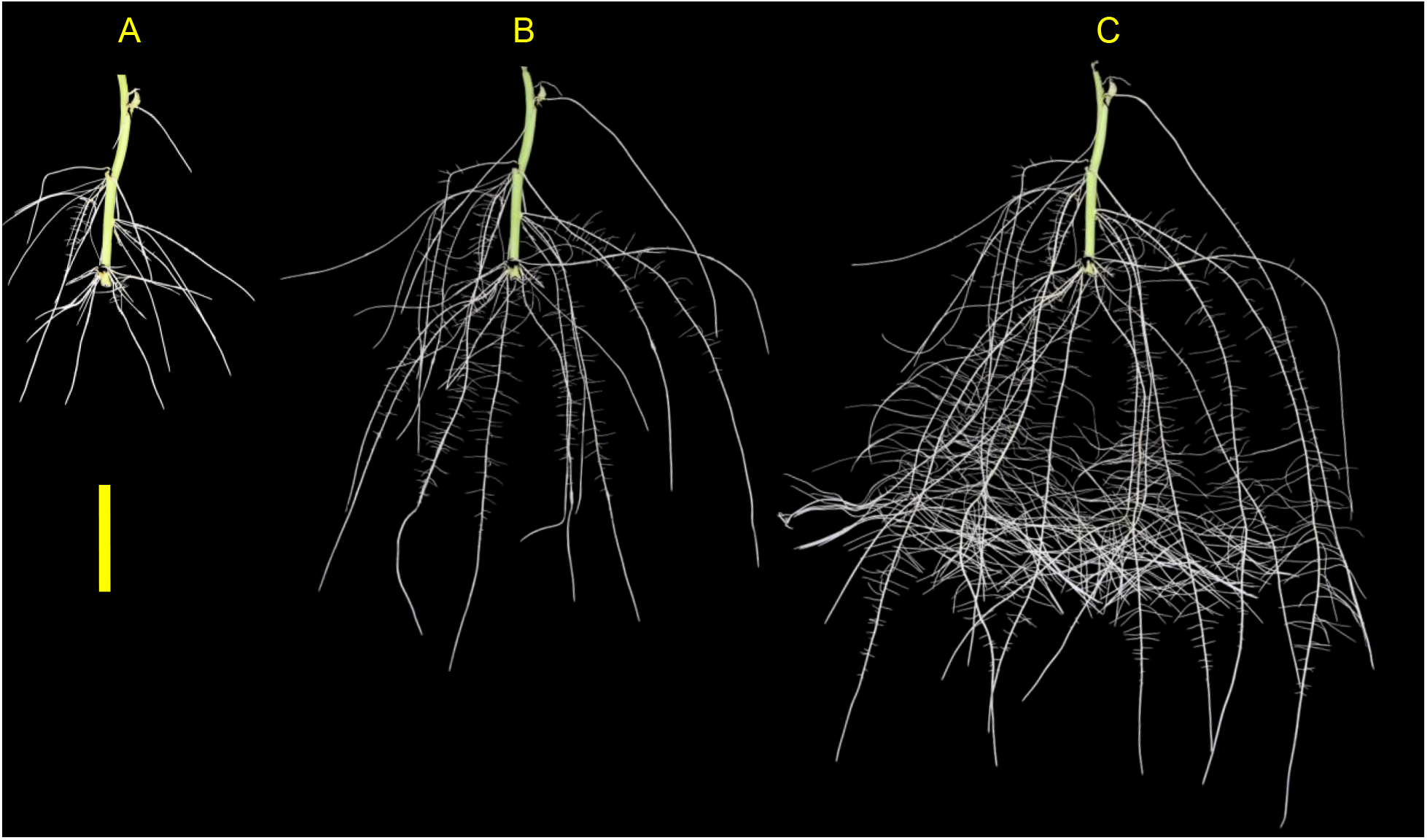
Example images showing root morphology and development of ‘NC10-275’ grown in the root phenotyping platform. Images were taken at (A) 5, (B) 10, and (C) 15 days after planting (scale bar = 10 cm).

### Root phenotype variation

Phenotyping of both sweetpotato genotypes produced root systems that were imaged and assessed every five days until day 20. Variations in several root traits between both genotypes were substantial. Significant variations were detected after day 10 in total root length (CV = 0.24 to 0.33, total root surface area (CV = 0.28 to 0.31), total root volume (CV = 0.07 to 0.32), and number of tips (CV = 0.22 to 0.48). After day 20, all root traits differed significantly for both genotypes (Table 2). Specifically, ‘NC10-275’ had a higher total root length, greater total surface root area, and larger total root volume compared to ‘Covington’. However, on average ‘Covington’ had a larger root diameter compared to ‘NC10-275’ on every sampling day (Table 2).

**Table 2.**
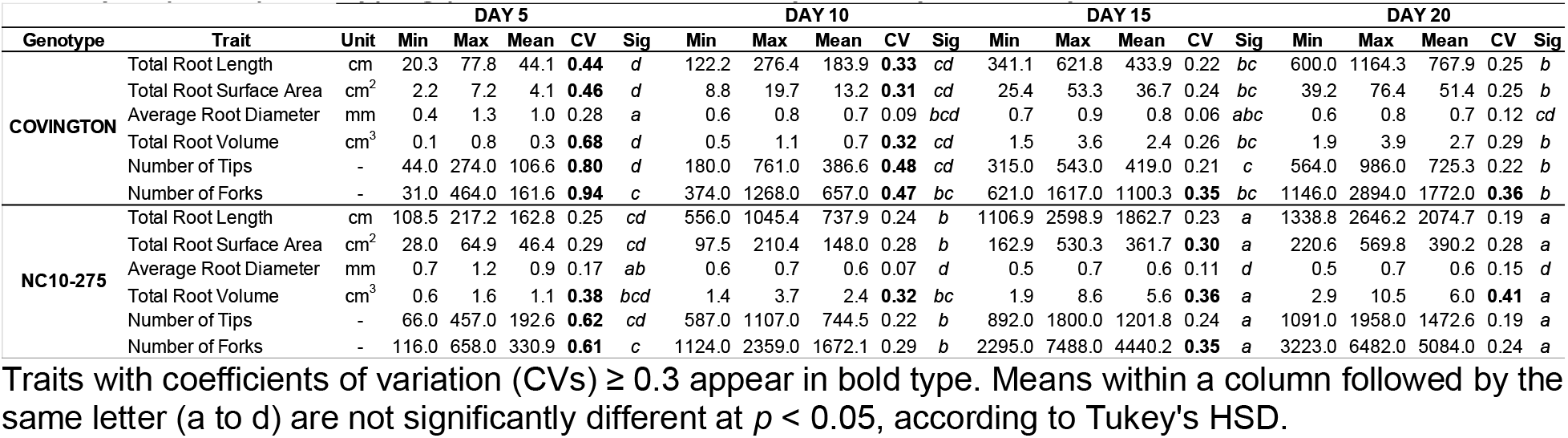
Descriptive statistics of six measured root traits in ‘Covington’ and ‘NC10-275’ grown in a 2D semi-hydroponic phenotyping platform assessed every five days until day 20.

| Genotype | Trait | Unit | DAY 5 |  |  |  |  | DAY 10 |  |  |  |  | DAY 15 |  |  |  |  | DAY 20 |  |  |  |  |
| --- | --- | --- | --- | --- | --- | --- | --- | --- | --- | --- | --- | --- | --- | --- | --- | --- | --- | --- | --- | --- | --- | --- |
|  |  |  | Min | Max | Mean | CV | Sig | Min | Max | Mean | CV | Sig | Min | Max | Mean | CV | Sig | Min | Max | Mean | CV | Sig |
| COVINGTON | Total Root Length | cm | 20.3 | 77.8 | 44.1 | **0.44** | d | 122.2 | 276.4 | 183.9 | **0.33** | cd | 341.1 | 621.8 | 433.9 | 0.22 | bc | 600.0 | 1164.3 | 767.9 | 0.25 | b |
|  | Total Root Surface Area | cm<sup>2</sup> | 2.2 | 7.2 | 4.1 | **0.46** | d | 8.8 | 19.7 | 13.2 | **0.31** | cd | 25.4 | 53.3 | 36.7 | 0.24 | bc | 39.2 | 76.4 | 51.4 | 0.25 | b |
|  | Average Root Diameter | mm | 0.4 | 1.3 | 1.0 | 0.28 | a | 0.6 | 0.8 | 0.7 | 0.09 | bcd | 0.7 | 0.9 | 0.8 | 0.06 | abc | 0.6 | 0.8 | 0.7 | 0.12 | cd |
|  | Total Root Volume | cm<sup>3</sup> | 0.1 | 0.8 | 0.3 | **0.68** | d | 0.5 | 1.1 | 0.7 | **0.32** | cd | 1.5 | 3.6 | 2.4 | 0.26 | bc | 1.9 | 3.9 | 2.7 | 0.29 | b |
|  | Number of Tips | - | 44.0 | 274.0 | 106.6 | **0.80** | d | 180.0 | 761.0 | 386.6 | **0.48** | cd | 315.0 | 543.0 | 419.0 | 0.21 | c | 564.0 | 986.0 | 725.3 | 0.22 | b |
|  | Number of Forks | - | 31.0 | 464.0 | 161.6 | **0.94** | c | 374.0 | 1268.0 | 657.0 | **0.47** | bc | 621.0 | 1617.0 | 1100.3 | **0.35** | bc | 1146.0 | 2894.0 | 1772.0 | **0.36** | b |
| NC10-275 | Total Root Length | cm | 108.5 | 217.2 | 162.8 | 0.25 | cd | 556.0 | 1045.4 | 737.9 | 0.24 | b | 1106.9 | 2598.9 | 1862.7 | 0.23 | a | 1338.8 | 2646.2 | 2074.7 | 0.19 | a |
|  | Total Root Surface Area | cm<sup>2</sup> | 28.0 | 64.9 | 46.4 | 0.29 | cd | 97.5 | 210.4 | 148.0 | 0.28 | b | 162.9 | 530.3 | 361.7 | **0.30** | a | 220.6 | 569.8 | 390.2 | 0.28 | a |
|  | Average Root Diameter | mm | 0.7 | 1.2 | 0.9 | 0.17 | ab | 0.6 | 0.7 | 0.6 | 0.07 | d | 0.5 | 0.7 | 0.6 | 0.11 | d | 0.5 | 0.7 | 0.6 | 0.15 | d |
|  | Total Root Volume | cm<sup>3</sup> | 0.6 | 1.6 | 1.1 | **0.38** | bcd | 1.4 | 3.7 | 2.4 | **0.32** | bc | 1.9 | 8.6 | 5.6 | **0.36** | a | 2.9 | 10.5 | 6.0 | **0.41** | a |
|  | Number of Tips | - | 66.0 | 457.0 | 192.6 | **0.62** | cd | 587.0 | 1107.0 | 744.5 | 0.22 | b | 892.0 | 1800.0 | 1201.8 | 0.24 | a | 1091.0 | 1958.0 | 1472.6 | 0.19 | a |
|  | Number of Forks | - | 116.0 | 658.0 | 330.9 | **0.61** | c | 1124.0 | 2359.0 | 1672.1 | 0.29 | b | 2295.0 | 7488.0 | 4440.2 | **0.35** | a | 3223.0 | 6482.0 | 5084.0 | 0.24 | a |

Traits with coefficients of variation (CVs) $\geq 0.3$ appear in bold type. Means within a column followed by the same letter (a to d) are not significantly different at $p < 0.05$ , according to Tukey's HSD.

### Root Length Pattern and Root-Depth Index

Root length by depth between ‘Covington’ and ‘NC10-275’ was non-significant from 0 to 30 cm, however significantly different from 30 to 50 cm by day 20 (Figure 7). Specifically, ‘Covington’ displayed the largest portion of root length from 20-30 cm (37.3%), followed by 30-40 cm (26.2%) and 10-20 cm (25.4%). Whereas ‘NC10-275’ exhibited the largest portion of root length at 30-40 cm (32.2%) followed by 40-50 cm (29.8%) and 20-20 cm (15.8%). Lastly, ‘NC10-275’ showed a total higher root-depth index (35.4%) compared to ‘Covington’ (23.6%).

**Figure 7.**
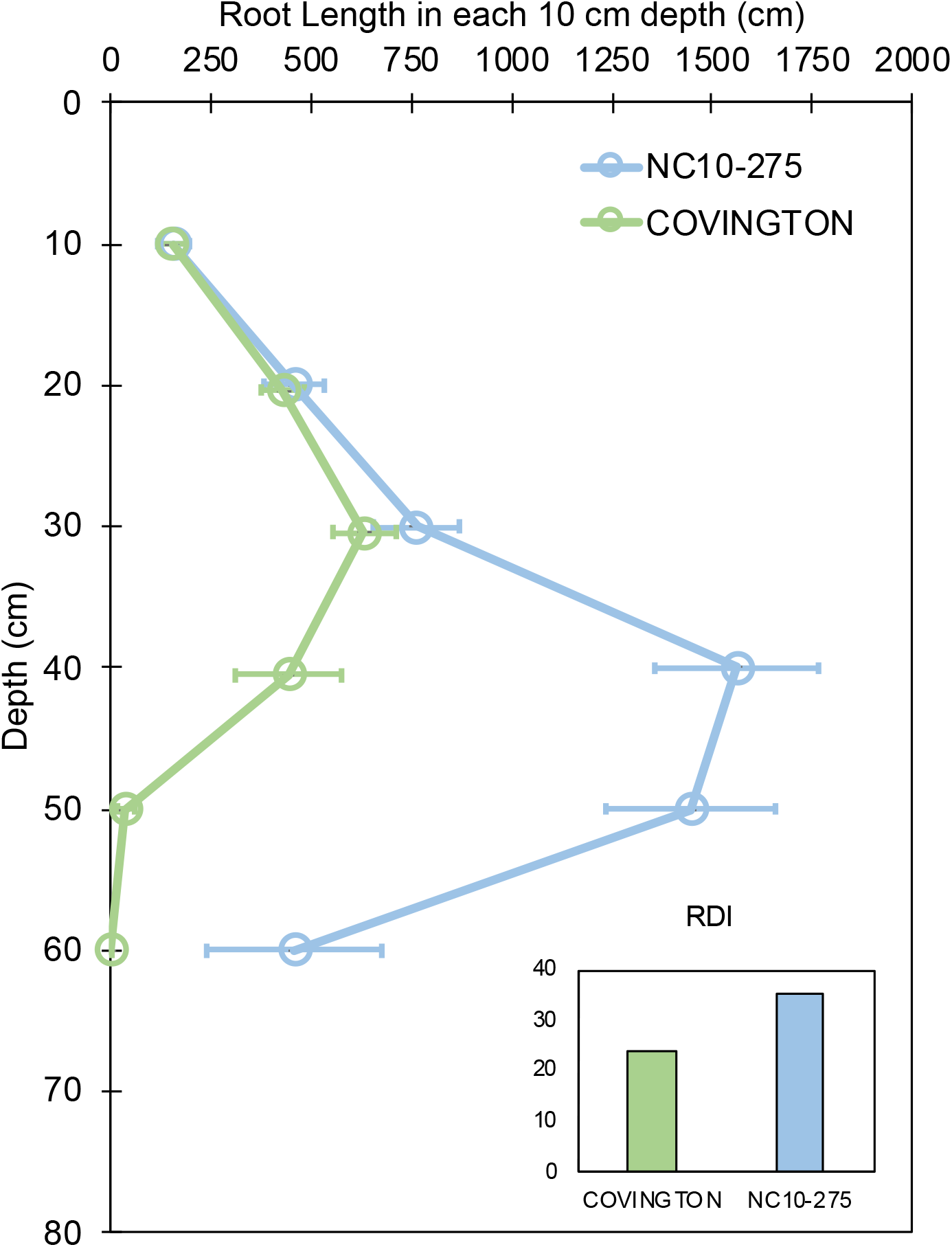
Root length in each 10 cm depth by soil depth and root depth index of ‘Covington’ and ‘NC10-275’ sampled at day 20. Data shown are means ± SE of 12 replicates of the two genotypes.

### Root Branching Density

Both genotypes selected for this study showed different lateral root branching density phenotypes when compared at each sampling day (Figure 8). Under the phenotyping platform, ‘Covington’ displayed a lower lateral root branching density compared to ‘NC10-275’ at day 5, however, this trend changed by day 10 until the end of the experiment. By day 10, both ‘Covington’ and ‘NC10-275’ presented similar lateral root branching density quantities, although by day 15 and 20, ‘Covington’ had significantly greater lateral root branching density when compared to ‘NC10-275’.

**Figure 8.**
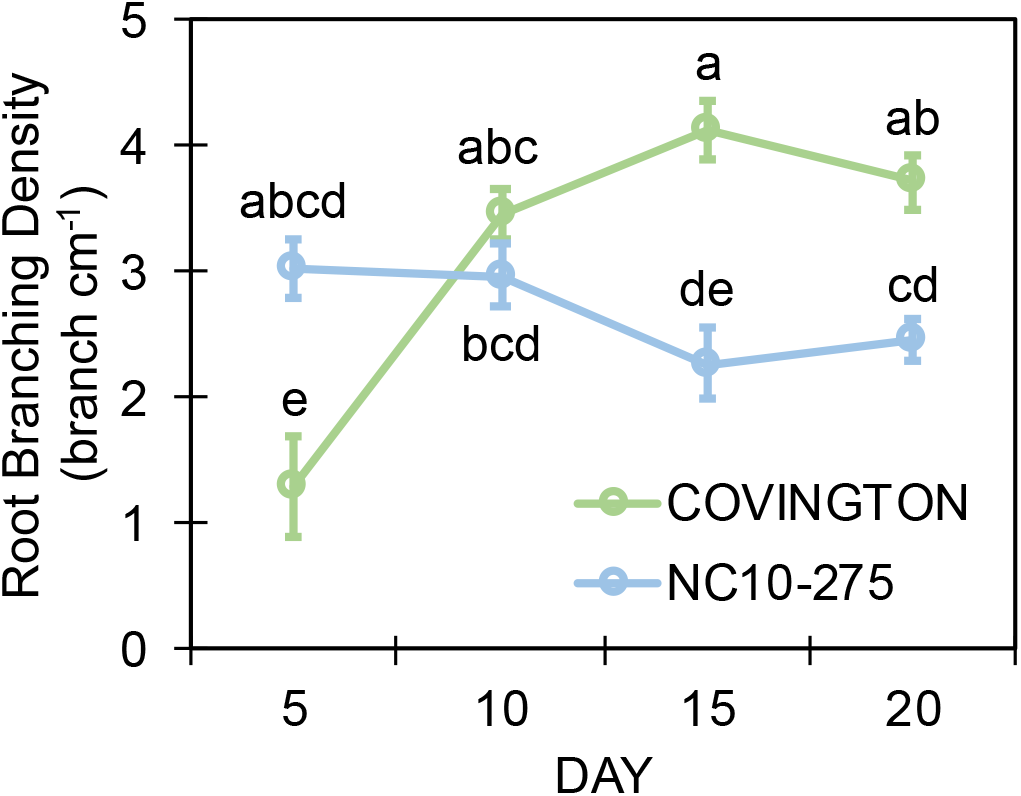
Lateral root branching density of adventitious roots at each sampling day for ‘Covington’ and ‘NC10-275’. Data shown are means of 12 replicates of the two genotypes in each sampling day. Different letters (a to d) represent significant differences at *p* < 0.05, according to Tukey’s HSD

### Root Length Distribution by Diameter Class

The total root length was divided into four diameter classes: very fine (0-0.6 mm), fine (>0.6 to ≤1.2 mm), large (>1.2 to ≤1.8 mm), and very large (>1.8 mm). In general, the root length distribution by diameter class of ‘NC10-275’ was significantly larger when compared to ‘Covington’ at each sampling day (Figure 9A and 9B). Specifically, the very fine and fine root length of ‘NC10-275’ was greater than that of ‘Covington’ at day 5 (very fine: 56.9 cm compared to 18.9 cm; fine: 95.3 cm compared to 14.5 cm), day 10 (very fine: 427.3 cm compared to 94 cm; fine: 312.4 cm compared to 78.3 cm), day 15 (very fine: 995.2 cm compared to 180.7 cm; fine: 531.4 cm compared to 157.9), and day 20 (very fine: 1184.9 cm compared to 429.7 cm; fine: 541.5 cm compared to 240.9 cm) respectively. Overall, the very fine and fine root diameter classes represented the longest root length for both ‘NC10-275’ and ‘Covington’ (97.7% and 92.4% of the total length) respectively. There was small to no significant differences in root length distribution for large to very large root diameters between genotypes (Figure 9A and 9B).

**Figure 9.**
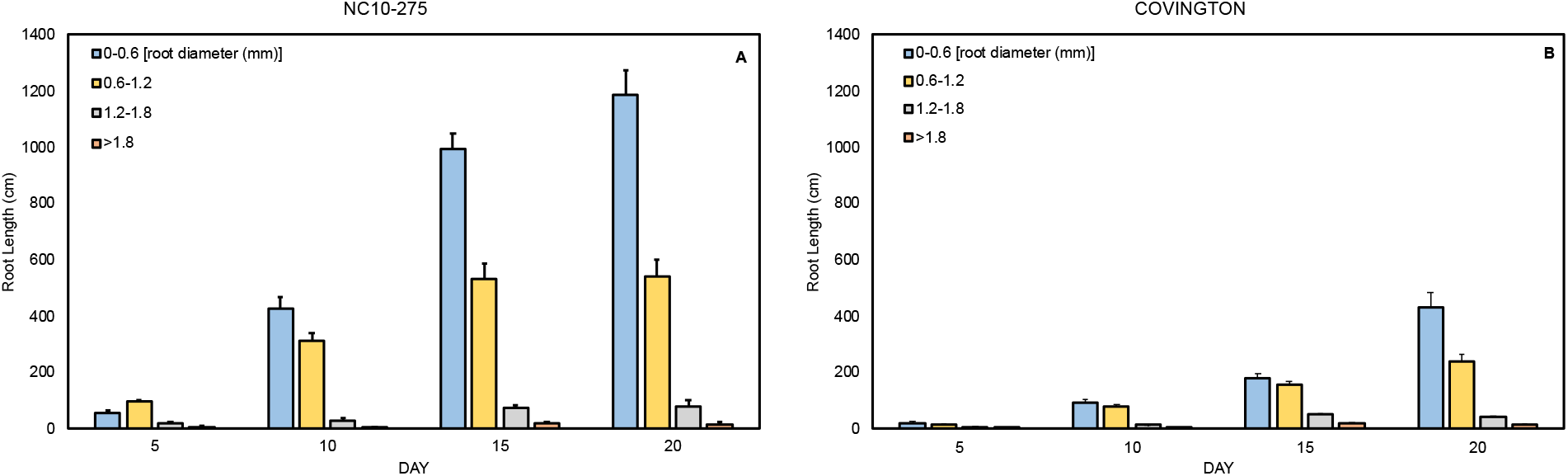
Mean root length distribution by diameter class of (A) ‘NC10-275’ and (B) ‘Covington’ at each sampling day. Data shown are means of 12 replicates of the two genotypes in each sampling day.

## Discussion

The main reasons for developing this study were two-fold: first, to development a cheap, cost-effective, and efficient phenotyping system for examining stem-derived (i.e., adventitious/nodal) roots and second, to examine the inherent genetic variations in root traits among a commercially available sweetpotato clone and an unreleased breeder line. The combination of these two approaches could provide the basis for future root models and facilitate three-dimensional root architecture for selecting superior root traits for sweetpotato breeding programs worldwide. This experiment using sweetpotato was devised to test the efficiency of the phenotyping system and the performance of cut and prepared ‘slips’ in the system. The results of this pilot study will provide information for future follow-up screening experiments using the same plant species. The rationale for the use of sweetpotato, considered a root and tuber crop (RTC) model organism is that this root crop species (as well as other RTCs), have lingered behind the well-studied “model” crop species like maize, rice, soybean, and wheat, where the knowledge of RSA has already led to considerable advances in the ability of these crops to exploit soil resources under low-input conditions.

The root growth phenotyping platform described here conforms to a ‘soilless 2D root phenotyping platform’ that uses a semi-hydroponic medium similar to many published reports (Chen *et al*., 2022; Adu *et al*., 2014; Chen *et al*., 2011; Le Marie *et al*., 2014). Specifically, the phenotyping platform allowed for a clear visualization of sweetpotato stem-derived adventitious roots growing on each of the individual growth units. In addition, the platform allowed for the growing of eight ‘slips’ simultaneously that could be removed individually every five days for imaging. The root growth phenotyping platform is comparable to the ‘pouch-and-wick’ system that allows for an *in situ* observation of adventitious roots based on germination paper. We developed this system because it is affordable, expandable, simple to operate, and can be used to evaluate early RSA with high efficacy. Also, as the system is expandable, it can conform to increased repetitions if necessary. However, attention is needed when removing the individual growth units for imaging as sweetpotato’s root system are fragile and root damage may occur. During root imaging, each individual growth unit was maintained moist and exposure time minimal to avoid roots from drying out. Per our observations, the root growth phenotyping platform is both semi-hydroponic and semi-aeroponic, which builds on the advantages of a strict hydroponic or aeroponic system. Since the root growth phenotyping platform uses an external and independent irrigation system (ebb-and-flow system) connected to each root growth unit, nutrient solutions can be prepared and re-stocked minimizing disruptions to the root growth unit. That said, this root growth phenotyping platform has the potential for root plasticity studies under water and nutrient stress. To our knowledge, is the first report of an *in situ* soilless 2-D root growth phenotyping system used on sweetpotato ‘slips’.

‘NC10-275’ is an unreleased breeder line that is considered “drought or wilt tolerant” and mainly used by the Sweetpotato Breeding and Genetics Program at North Carolina State University as a parental line to exploit its abiotic stress tolerance for breeding ornamental sweetpotato (*pers. comm*), whereas ‘Covington” is one of the most important commercial sweetpotato grown in the United States characterized by high yields and quality. The root systems of both ‘Covington’ and ‘NC10-275’ revealed unique root morphological features and root traits when developed in the root growth phenotyping platform. The root system of each genotype maintained comparable growth patterns until after day 5. We compared images and data from each sampling day between both genotypes and determined that after day 20 both root systems presented a higher diversity of root traits compared earlier sampling days. Differences in root traits after day 10 included root length, surface root area, root diameter, root volume, root depth, and root branching density. For example, it was noted that ‘NC10-275’ grew at a faster pace compared to ‘Covington’ increasing the abovementioned traits in favor of ‘NC10-275’. However, average root diameter and root branching density was higher in ‘Covington’. Also, it is noteworthy that the root length distribution by diameter class of ‘NC10-275’ was greater in all instanced measured. Root length pattern was increased only after 30 cm depth (measured at day 20) for both genotypes, yet after 30 cm depth ‘NC10-275’ expanded its root length exceeding that of ‘Covington’. Taken as a whole, all root traits measured revealed contrasting differences between both genotypes examined. ‘NC10-275’ exhibited an earlier root growth habit with more deeply distributed root system than ‘Covington”, probably due to its inherent drought tolerance where roots present a more vertical growth pattern. In contrast, ‘Covington’ revealed an overall reduced root volume, higher root branching density, and larger average diameter roots. This phenomenon could be explained from the basis on current agricultural management practices where fertilizer and water supply are abundant lessening the burden of root exploration for soil nutrients and water and investing more resources in storage root formation and swelling. These results could be confirmed by the higher root depth index (RDI) of the ‘NC10-275’, echoing the deeper root system of this genotype. Though root spreading (root width growth) was not accounted for, together with root depth pattern and root depth index are key traits for soil exploration for improving the acquisition of limiting resources. Regardless of both genotypes belonging to the same species (*I. batatas*), these results ratify the contrast that can be found between sweetpotato root systems. It is notable that no previous studies on the RSA or root traits of both ‘Covington’ and NC10-275 have been published. Nevertheless, the first published report measuring lateral root branching in sweetpotato was in 1949 (Koshimizu and Nishida, 1949), followed by other published reports years later (Villordon *et al*., 2012; Pardales and Yamauchi, 2003).

Though this research did not account for specific abiotic stress treatments [e.g., nitrogen (N), phosphorus (P), potassium (K) deficiencies, and/or water stress)] within the root growth phenotyping platform, there are now several published reports on the effects of N, P, K, and B deficiencies on RSA and root traits using mesocosms filled with sand or other substrates. For example, Villordon et al. (2013) demonstrated that lateral root branching jointly measured as lateral root length, number of lateral roots and lateral root density in ‘Beauregard’ was altered in response to variation in overall available N. Also, Villordon et al. (2020) revealed the existence of genetic variation for inorganic P efficiency in ‘Bayou Belle’, ‘Beauregard’ and ‘Orleans’ sweetpotato cultivars by measuring root lateral root number and lateral root density and found that these two root traits have been indirectly selected for inbreeding programs that focus on early storage root formation and stable yields across environments. Furthermore, Liu et al. (2017) showed differences in root length, surface area, root volume and average root diameter under controlled K and deficient K using two cultivars, Ningzishu 1 (sensitive to K deficiency) and Xushu 32 (tolerant to K deficiency). These results suggest potential genotypic differences in RSA and K absorption ability under K deficiency. Likewise, Wang et al. (2017), showed that increased K improved total root length, average root diameter and significantly increased the differentiation from adventitious roots to fibrous roots and tuberous roots. These root traits coupled with additional K could be beneficial to the increased number of storage roots per plant, early formation of storage roots, root biomass, and overall yield. Under differing B availability, Villordon and Gregorie (2021) showed evidence of cultivar-specific responses for reduced lateral root length, root length, and reduced storage root swelling in ‘Beauregard’, ‘Murasaki’, and ‘Okinawa’ cultivars.

## Conclusions

Root growth patterns for both genotypes tested retained comparable growth patterns until after five days in the phenotyping platform. After 20 days in the phenotyping platform both root systems showed the highest diversity and difference of root traits compared to earlier sampling days. Root length, surface root area, root volume, and root depth were higher in ‘NC10-275’. Average root diameter and root branching density were higher in ‘Covington’. Sweetpotato is a clonally propagated crop, sexual seeds are not used for planting, hence the experiment was performed with ‘slips’, the central unit of sweetpotato planting material used routinely in the field. In summary, this is the first report of a phenotyping system that uses a stem and not a sexual seed as starting material. This experiment confirmed genotypic variations in the early root system growth of sweetpotato using an ebb and flow soilless phenotyping platform. This phenotyping study was reproducible across the whole growing period and for both genotypes tested. However, one of the potential drawbacks of this system is the early inference of the potential performance of these genotypes in the field. Thus, under changing growing environments, roots may present specific responses making their inherent phenotypic plasticity critical for mining edaphic resources (Lynch *et al*., 2021). Yet, it is still possible to extrapolate early genotypic differences between sweetpotato germplasm and phenotypic plasticity under imposed stress treatments.

## Acknowledgements

The corresponding author would like to thank Dr. Jonathan Lynch (Penn State University) and Dr. Craig Yencho (North Carolina State University) for guidance.

## Notes

### Competing Interest Statement

The authors have declared no competing interest.

